# Oscillatory infrasonic modulation of the cochlear amplifier by selective attention

**DOI:** 10.1101/403865

**Authors:** Constantino D. Dragicevic, Bruno Marcenaro, Marcela Navarrete, Luis Robles, Paul H. Delano

## Abstract

Evidence shows that selective attention to visual stimuli modulates the gain of cochlear responses, probably through auditory-cortex descending pathways. At the cerebral cortex level, amplitude and phase changes of neural oscillations have been proposed as a correlate of selective attention. However, whether sensory receptors are also influenced by the oscillatory network during attention tasks remains unknown. Here, we searched for oscillatory attention-related activity at the cochlear receptor in humans. We used an alternating visual/auditory selective attention task and measured electroencephalographic activity simultaneously to distortion product otoacoustic emissions (a measure of cochlear receptor-cell activity). In order to search for cochlear oscillatory activity, the otoacoustic emission signal, was included as an additional channel in the electroencephalogram analyses. This method allowed us to study dynamic changes of cochlear oscillations in the same range of frequencies (1-35 Hz) in which cognitive effects are commonly observed in electroencephalogram works. We found the presence of low frequency (<10 Hz) brain and cochlear amplifier oscillations during periods of selective attention to visual and auditory stimuli. Notably, switching between auditory and visual attention modulates the amplitude and the temporal order of brain and inner ear oscillations. These results extend the role of the oscillatory activity network during cognition in neural systems to the receptor level.

## Introduction

In natural environments animals are surrounded by a great number of sensory stimuli. As the nervous system has a limited capacity for processing all sensory stimuli, individuals require of attention to focus their cognitive resources on important stimuli. Selective attention is a top-down form of attention in which one sensory modality is important to accomplish a given task and the other modalities are irrelevant or even distracting [1]. At the mechanistic level, it has been proposed that selective attention can function as a biological filter, meaning that neural responses to the attended stimulus are enhanced, while responses to unattended stimuli can be diminished. Whether these processes occur only at the central nervous system or also at more peripheral levels has remained controversial for many [2-4].

Evidence in mammals demonstrates that in the case of visual selective attention with auditory distractors, a modulation of the gain of auditory responses is clearly seen at the cortical level [3,5], while at the peripheral level, conflicting results have been reported, including positive [6-8] and negative findings [3,4,9]. However, the modulation of the gain of sensory responses by attention does not explain all possible brain mechanisms of top-down attention. Another proposed mechanism for attention arises from the known oscillatory nature of the nervous system, which has been suggested as a general mechanism for perception and cognition in vertebrate and invertebrate animals [10-12]. Amplitude changes in specific frequency bands or the entrainment of neuronal oscillations have been proposed as mechanisms of attentional selection [13-15], which could allow large or local scale synchronization among different brain areas [10,16]. However, whether cortical oscillations during selective attention to visual stimuli modulate cochlear responses at the receptor level is unknown. Here, we used an alternating visual/auditory selective attention task in humans (based on [17]) and measured electroencephalographic (EEG) activity simultaneously to a virtual channel of the amplitude of distortion product otoacoustic emissions (DPOAE) that allowed us to examine in the frequency domain of single-trials, the dynamics between cortical electrical oscillations and hypothetical oscillatory activity of the cochlear amplifier [18,19].

## Results

Continuous 32-channel EEG and DPOAE were recorded in 14 subjects performing alternating tasks that required attentional switches between visual and auditory perceptual modalities (Fig 1). Both modalities required high temporal acuity in detecting time in a revolving clock (visual) or a brief gap of silence embedded in continuous DPOAE-eliciting pairs of tones (auditory). Time and frequency averaged EEG and DPOAE signals were analyzed in a period of delimited high expectancy (selective attention) in both modalities, corresponding to the epochs before the appearance of auditory and visual targets (from 0 ms to 1500 ms) and were compared with the previous period (−1500 ms to 0 ms). The DPOAE channel was calculated using the amplitude of the frequency band surrounding (±100 Hz) the 2f1-f2 component (Supplementary Figs. 1 and 2). In order to analyze EEG and DPOAE signals as part of the same functional network during selective attention, we added the DPOAE amplitude signal as an additional channel in EEG analyses.

**Fig 1.**
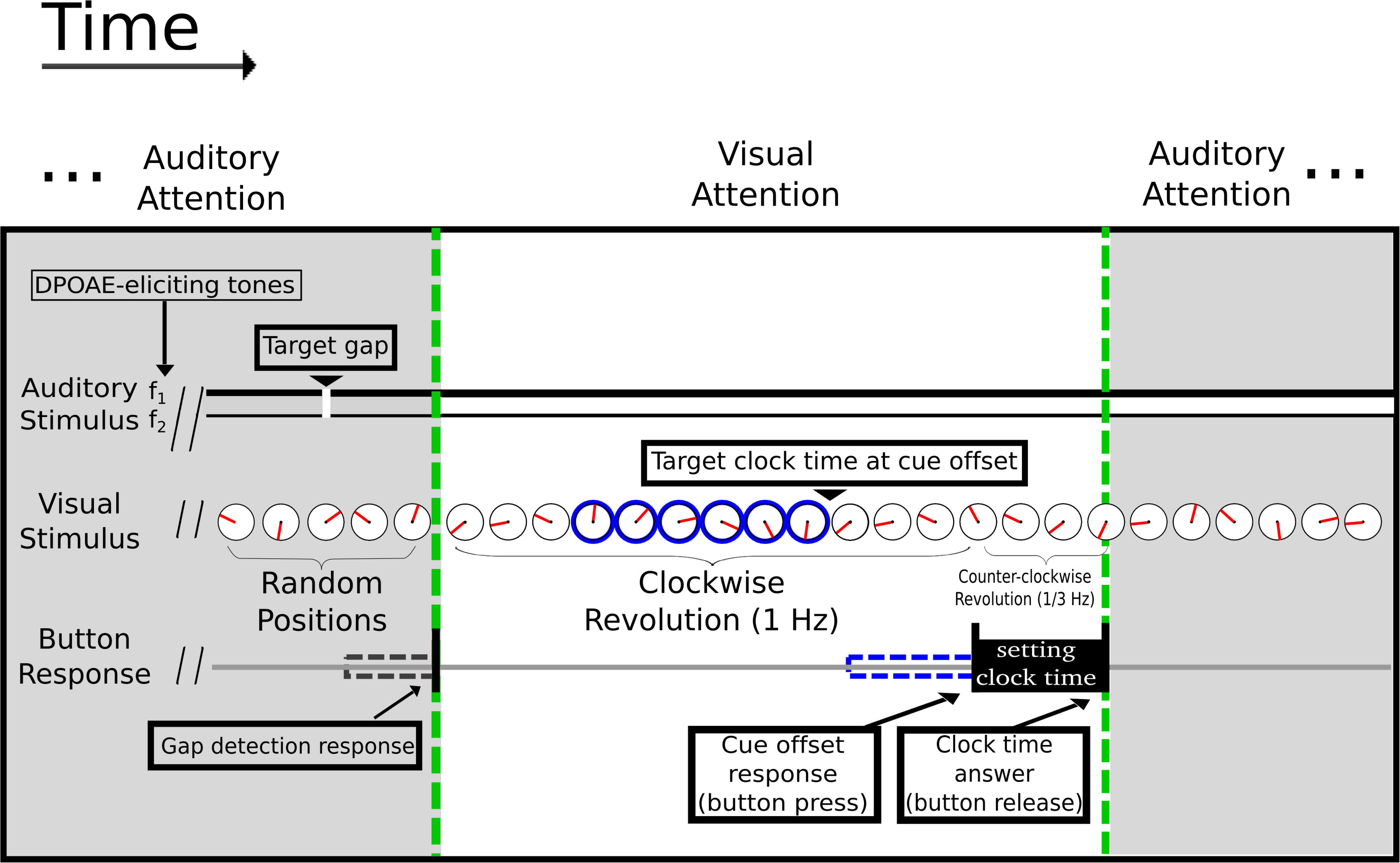
Illustration of the visual/auditory selective attention task. Subjects were required to alternate their attention between the visual and auditory modalities after each trial. The green dashed lines delimit the periods of auditory (grey shaded area) and visual (white area) attention. During visual attention a minute indicator (shown in red) rotates clock-wise at 1 Hz. After a random passive period of 2,000-2,500 ms, a peripheral clock rim appears as a temporal visual cue (shown in blue) and remains on for a variable period of 1,500-2,500 ms. Subjects were asked to report the position of the clock at the off-set of the peripheral clock rim. Simultaneously, in order to evoke DPOAEs, two tones (f1 and f2) were presented continuously during the visual attention task. On selective auditory attention trials (shaded grey), volunteers were required to report a brief (2-4 ms) silence gap embedded in the continuous DPOAE-eliciting tones. Gap detection triggered the switch to the visual task.

**Fig 2.**
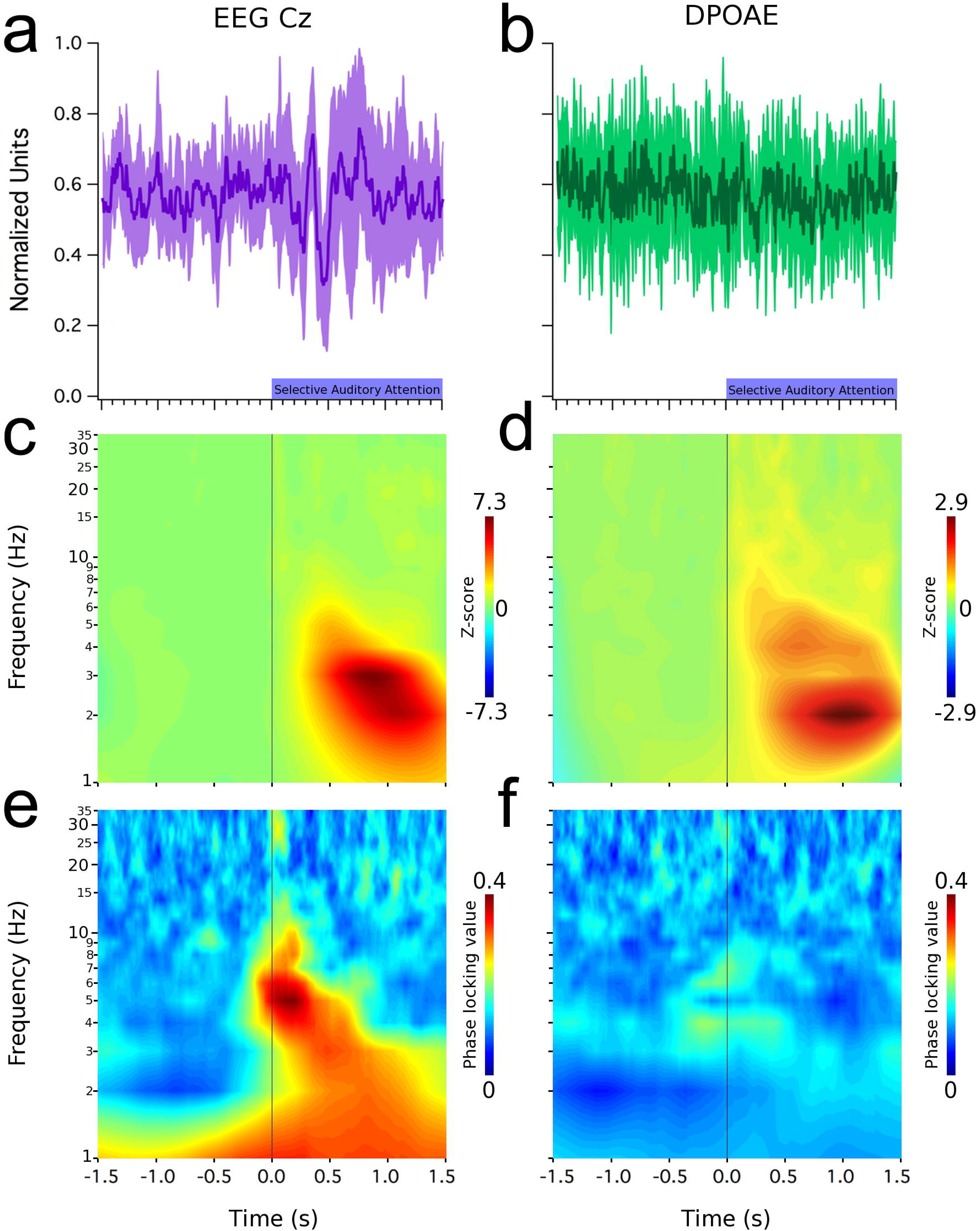
Low-frequency oscillations (< 10 Hz) in EEG (Cz electrode) and in the cochlea (DPOAE) during selective auditory attention. Time and frequency graphs represent grand average data. Left column shows: (a) evoked oscillations at Cz electrode, (c) time spectrum and (e) an increased phase locking value of these brain oscillations. Right column shows: (b) a lack of evoked effects in the amplitude of DPOAEs, while (d) low frequency induced DPOAE amplitude oscillations can be observed between 1 and 7 Hz. (f) Notice the lack of phase lock in cochlear oscillations. Shaded purple and green areas in (a) and (b) represent data dispersion (± 1 standard deviation).

During the period of selective auditory attention, in which subjects had high expectancy for a silence gap embedded in continuous primary f1 and f2 tones that evoke DPOAE, an evoked potential appeared in the grand average of the EEG signal at Cz (Fig 2a), while in the same period a subtle non-significant reduction was observed in the DPOAE signal (Fig 2b). During this period, we also found the presence of low frequency oscillations (<10 Hz) in the brain (Fig 2c, EEG) and cochlear receptor (Fig 2d, DPOAE). EEG Cz oscillations were phase-locked to the onset of the auditory attention period (Fig 2e), while cochlear oscillations had enough jitter to show a complete absence of phase-locking (Fig 2f). Fig. 3 shows grand average results for the case of visual selective attention. A visual evoked response was clearly seen in the occipital EEG channels (Fig 3a), while the averaged DPAOE signal showed no effect (Fig 3b). Similarly to the auditory attention trials, the frequency analyses of EEG and DPOAE signals yielded the presence of low-frequency (<10 Hz) oscillations at the brain and cochlear levels (Fig 3c and 3d). The phase-locking values of these oscillations, show that the EEG signal coming from the occipital cortex is synchronized to the onset of the visual attention period (Fig 3e), while no phase-locking was seen at the cochlear level (Fig 3f).

**Fig 3.**
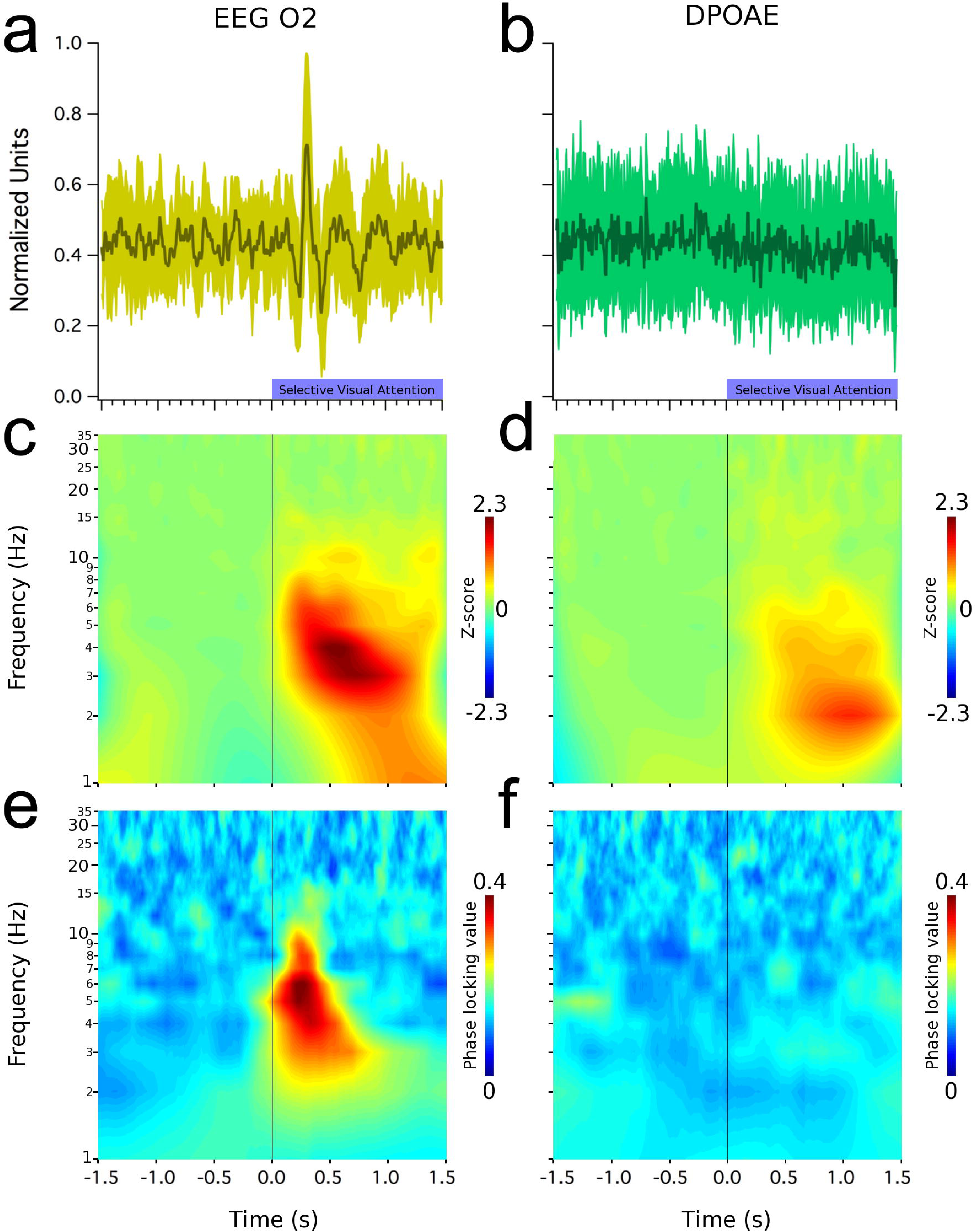
Low-frequency oscillations (< 10 Hz) in EEG (O2 electrode) and in the cochlea (DPOAE) during selective visual attention. All graphs represent grand average data. Left column shows evoked oscillations at the right occipital cortex (a), time spectrum (c) and (e) increased phase locking values of these brain oscillations. Right column shows (b) a lack of evoked effects in DPOAE amplitudes, while (d) low frequency induced oscillations can be observed between 1 and 7 Hz. (f) Notice the lack of phase lock in cochlear oscillations. Shaded green areas in (a) and (b) represent data dispersion (± 1 standard deviation).

In order to compare amplitudes and temporal dynamics of single trial EEG and cochlear oscillations (Fig 4a and 4b), during visual and auditory attention, oscillation amplitudes (frequency band 1-7 Hz) were normalized as z-scores for both types of attention. For cochlear oscillations, comparisons showed a significant reduction in amplitude during periods of visual attention (0.97 ± 0.48 z) compared with those of auditory attention (1.63 ± 1.15 z) (Z_(20)_= -2.089, p=0.038, U-Mann-Whitney). Regarding the temporal relation between brain and cochlear oscillations, we analyzed the half-time necessary to get 50% of the normalized amplitude of EEG and DPOAE oscillations (1-7 Hz) during visual and auditory attention. During visual attention occipital cortex oscillations preceded DPOAE oscillations (Fig. 4c), while during auditory attention, this order was inverted as the cochlear amplifier oscillations preceded occipital cortex oscillations (Fig. 4d).

**Fig 4.**
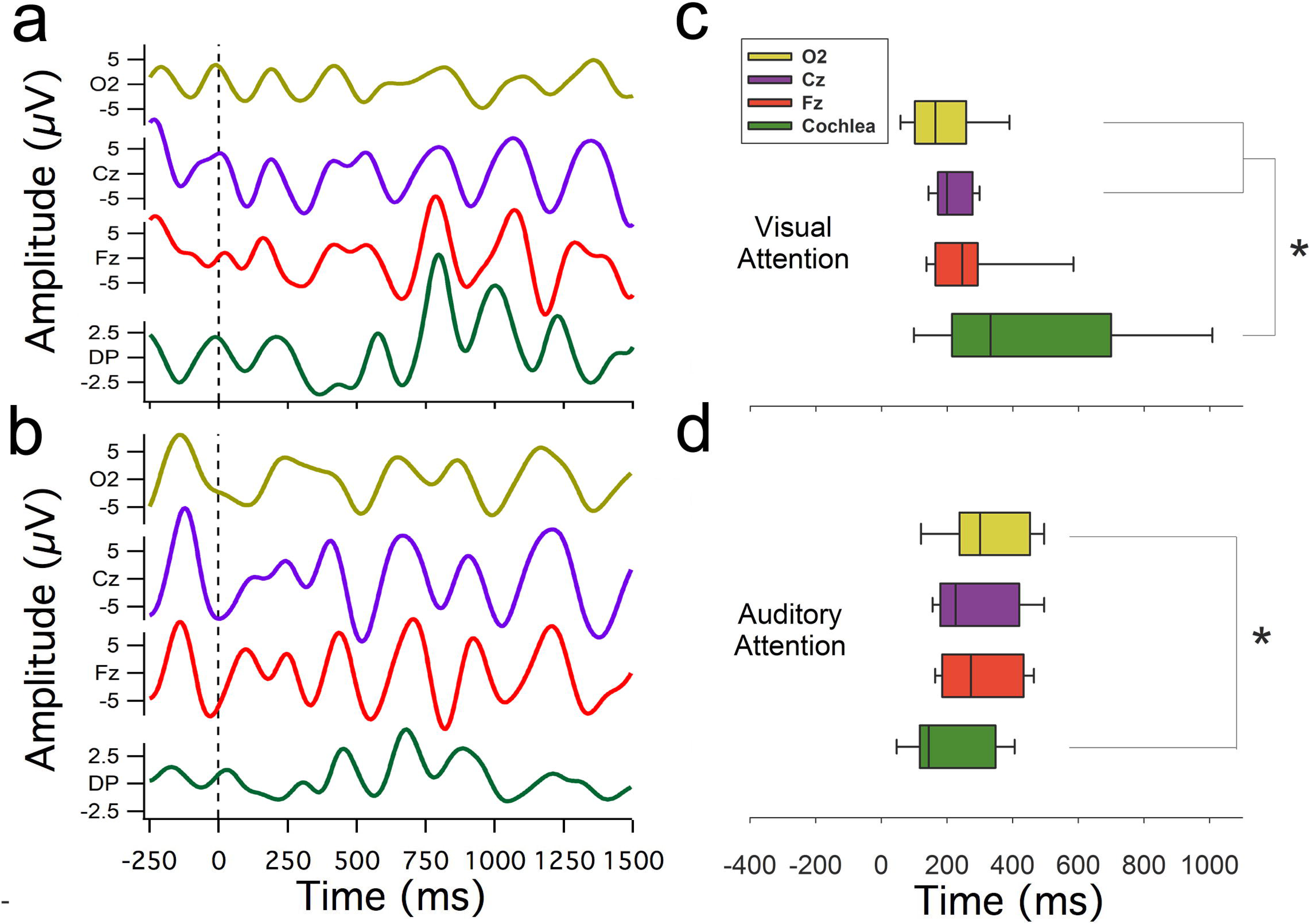
Temporal dynamics of brain and cochlear oscillatory activity. Single trial EEG and DPOAE oscillations filtered between 1 and 7 Hz are shown in (a) (visual attention) and (b) (auditory attention). Box plots in (c) and (d) represent, for the visual and auditory case respectively, the group distribution of the half times of the oscillatory peak activity in the 1-7 Hz frequency band of EEG recordings from O2 (yellow), Cz (purple), Fz (red) and cochlea (green, DPOAE signal). Asterisks denote a statistically significant difference of half-peak times (p<0.05, Mann-Whitney tests) between DPOAE amplitude and O2 for both modalities, and also with Cz in the visual attention case. Notice wider distribution of half times in DPOAE oscillations (green boxes) during visual attention as compared to auditory attention.

## Discussion

In the present work, we found the presence of low-frequency (<10 Hz) EEG and DPOAE oscillations during periods of selective attention to visual and auditory stimuli. These findings expand the framework of the oscillatory mechanisms in the attentional network, as many authors consider primary sensory cortices as the earliest brain regions that could be affected by these mechanisms, while the present work adds the cochlear receptor as the most peripheral structure modulated by the oscillatory network of attention.

It is important to remind that otoacoustic emissions are sounds –pressure waves-emitted by the inner ear that can be measured with a sensitive microphone positioned at the external ear canal [20]. They are thought to reflect the electro-motility of outer hair cells of the cochlear receptor, which is the proposed cellular mechanism of cochlear amplification [19,21,22]. The frequency band (<10 Hz) of the amplitude oscillations modulating the DPOAE (2f1-f2) that we found is located below the human audible range (which goes from 20 to 20,000 Hz), and therefore, these low-frequency oscillations can be considered as infrasound waves [23].

Whether attention modulates the cochlear receptor has remained controversial for many years [3,4]. A number of works have found top-down effects of visual attention at the cochlear amplifier measuring otoacoustic emission amplitudes [8,24-26], however other authors have failed to find them [3,4,9]. In our work, we also explored frequency specific oscillatory mechanisms for attentional selection [14] at the cochlear level. In the present work, although we did not find a modulation of the mean amplitude of evoked DPOAEs by selective attention (Figs 2b and 3b), we did find low-frequency oscillations in DPOAE amplitudes produced by selective attention.

A previous work [8] attempted to investigate the modulatory effects of selective attention on EEG and DPOAE recordings. Although they found a significant reduction of the power of alpha oscillations at the occipital cortex and a significant decrease of DPOAE amplitudes during visual attention, they did not find any statistical relation between both measures. In our case we found DPOAE amplitude oscillations with a frequency band similar to the EEG oscillations (1-7 Hz) during auditory (Fig 2) and visual (Fig 3) attention. Moreover, the temporal order of the electrical brain oscillations at the occipital cortex and the mechanical oscillations at the cochlear receptor was inverted depending on the attended modality (Fig 4c, d). If the subjects were attending to the visual task, then the low-frequency occipital oscillations preceded DPOAE amplitude oscillations (Fig 4c). On the contrary, if attention was directed to the acoustic stimuli, then cochlear oscillations led occipital cortex waves (Fig 4d). Thus, the temporal order between occipital cortex and DPOAE oscillations was inverted by switching between auditory and visual attention tasks.

The low-frequency oscillations that we observed in our tasks (<10 Hz) can be classified as delta and theta EEG oscillations [12]. Regarding theta oscillations, it has been theorized that in cognitive tasks in humans they emerge in the frontal cortex [27] and serve as a time reference for the dynamic assembly of different neural populations, (e.g. hippocampus), by increasing and decreasing the firing rate probability of single neurons subjected to the extracellular local field potentials induced by more global cortical oscillations [28,29]. In the context of attention, theta oscillations have been found in cross-modal paradigms involving visual and auditory attention [30,31]. These authors showed that theta allows fronto-parietal top-down modulation of visual and auditory cortices during cross-modal attention. In our work we extend the oscillatory network of top-down attention towards the cochlear receptor, showing that occipital EEG low-frequency oscillatory activity precedes mechanical oscillations in the cochlear amplifier during visual attention. We propose that the descending pathways from the auditory cortex to the cochlear receptor that comprise the auditory efferent system [32,33], are the most probable neural pathways that could explain the modulation of low-frequency oscillatory amplitude changes of DPOAE during selective attention to visual or auditory stimuli.

During visual attention cochlear oscillations have a significant temporal jitter as compared with EEG oscillations (Fig. 4c), which might reflect an active process to reduce the peripheral entrainment of auditory stimuli during visual attention. This possible mechanism would be in agreement with works showing that low-frequency oscillations can modify the mechanical sensitivity of the cochlear receptor [18,34]. On the other hand, during auditory attention, cochlear oscillations precede occipital EEG low-frequency oscillations, and significant less jitter is observed (Fig. 4d), thus allowing entrainment of cochlear responses to auditory stimuli. The latter proposal would be in agreement with a general mechanisms of oscillatory entrainment during attention to the corresponding relevant stimulus [14,35].

In summary, we found EEG and cochlear amplifier infrasonic oscillations during periods of visual and auditory attention. Moreover, the attentional switch between visual and auditory attention modulates the amplitude and the temporal order of brain and inner ear oscillations. These results extend the role of oscillatory activity in the nervous system during cognition to the receptor level.

## Materials and methods

### Ethics statement

The study was approved by the ethical committee from the Clinical Hospital of the Universidad de Chile, permission number: OAIC 016/20042016. All procedures were conducted in accordance to this approved protocol and to national regulations.

### Participants

Fourteen right-handed volunteers participated in our experiments (four females, mean age 24.2 ± 4.0 (SD, standard deviation)). All volunteers gave written consent and have no hearing and no neurological impairments. Because of the strict procedures to remove EEG and DPOAE artifacts (see below) we excluded electrophysiological data from one subject from the visual attention task, and five subjects from the auditory attention task.

### General experimental procedures

All procedures were carried out in an acoustically isolated room designed for audiological and electrophysiological evaluations inside the Clinical Hospital of Universidad de Chile. Electroencephalographic signals (32-channel EEG, Tucker Davis Technologies®) and continuous DPOAE dynamics were recorded simultaneously from each subject, which performed in the alternating tasks of visual and auditory selective attention, as depicted in Fig 1. A National Instruments® I/O card (model NI6321) together with a Tucker-Davis Technologies® multiprocessor (model RZ6) along with a desktop computer controlled the experiment as coded in Labwindows/CVI 2009 (C language) and System 3 (Tucker-Davis Technologies®) programming environments.

Before positioning any measuring device on subjects, we discarded earwax obstruction in both ear canals. Then we set up EEG recording, followed by fitting the insert earphones and microphone for DPOAE recording. Then we calibrated the intensity of sound delivery, and measured DPOAE at different frequencies (between 0.5 and 4 kHz) to choose the primary tones parameters that elicited the cleanest DPOAE signal. Only then we gave instructions to the subjects, verifying their understanding and execution through supervised training blocks, after which the main experiment began.

### DPOAE

During the experimental protocol, primary tones f1 and f2 were presented to the right ear continuously in order to elicit 2f1-f2 DPOAE, which were recorded continuously during approximately 8 minutes by a microphone (ER-2, Etymotic Research®) sealed in the external right ear canal. Before the experimental protocol, a set of nine pairs of tones with corresponding frequencies (f1 and f2) and intensities (L1 and L2) were generated with L2 fixed at 55 dB SPL and F2 logarithmically spaced between 1-5 kHz. Primary tone parameters selected by this procedure can be seen in Supplementary Fig. 1. Calibration of each frequency was done separately for both phones (phone A dealing with f1 and phone B with f2), by playing, adjusting, and replaying long tones (4,000 ms) to reach a constant sound pressure level of 50 dB SPL. Then we determined which pair of tones produced the largest DPOAE signals, based on 20 presentations of each tone pair (stimuli lasted for 1,000 ms and had an inter-stimulus interval of 500 ms). This was judged and manually selected by the experimenters based on both graphical inspection of the spectrum of the average DPOAE signal, and three measures for each tone pair: a) absolute peak amplitude at DPOAE frequency 2f1 – f2, b) surrounding noise amplitude of surrounding frequencies (± 10 Hz), and c) standard deviation of surrounding noise. With these parameters, we calculated the difference between DPOAE amplitude and surrounding noise (a – b), and the difference between this difference and the standard deviation of surrounding noise ([a – b] -c). The main experiment ran along with the tone parameters selected in this step.

### EEG

32 EEG channels referenced to the right earlobe and two bipolar electro-oculogram (EOG) recordings (for vertical and horizontal eye movements) were preamplified and digitized by battery powered Tucker-Davis Technologies® devices (PZ3 for EEG and RA4PA for EOG). Ring shaped Ag/AgCl electrodes and headcaps manufactured by EasyCap® were utilized. The elastic headcap (size 56 or 58) was secured with velcro under the chin. EEG electrodes positions complied with the 10-20 the standard system. Ground electrode was positioned on Fz. Electroconductive gel was applied to the scalp contacts posterior to cleansing with alcohol to keep impedances under 5 kΩ. A digital 0.1-100 Hz band-pass filter was applied along with a notch filter at 50 Hz. The output of this filtered data was saved with a sampling rate of 1 kHz.

### Attention tasks

During the visual attention tasks, f1 and f2 tones served as distractors. In contrast, during the auditory attention task the tones had to be attended for detecting a brief gap of silence (5 ms squared cosine ramps and 2-4 ms of complete silence). The perceptual modality to be attended alternated after valid responses or after the end of response time windows, 100-1,000 ms from target. Subject responses were given with the right thumb through a custom made push button. Subjects completed four experimental blocks of 44 trials for each modality, after at least 1 training block (explained below). Each block had an approximated duration of eight minutes.

### Visual task, stimuli and apparatus

The visual task starts with a passive (no attention) pseudo-random period of 2,000-2,500 ms. During this period, subjects are only required to maintain fixation at the center of a single-handed clock 4° in diameter. The clock hand revolves clockwise at 1 Hz, passing through each of 100 divisions or tick marks. To accomplish synchronization between custom software and the high refresh rate (100 Hz) monitor (Samsung® LED 23” 3D S23A700D), a time counter of the National Instruments® I/O card was configured to trigger screen refresh at the same 100 Hz. This ensures a smooth and coherent motion perception. At some point during the 2,000-2,500 ms passive period, a visual cue appears as a change in color of the external rim of the clock, indicating the period of visual selective attention (shown in blue on Fig 1). Subjects had to report immediately and as precisely as possible the clock hand position at the time of visual cue offset, occurring 1,500-2,500 ms from its onset. This task was adapted from a similar version implemented by other investigators [17]. A quick reaction was encouraged, but without interfering with response precision. To indicate the time at which subjects thought that cue offset occurred, the button had to be maintained pressed, inverting the rotation of the clock hand (from clockwise to counter-clockwise) and slowing down to 0.33 Hz, eventually passing over target position, where the button had to be released to set the response. No feedback was given about performance. Immediately after button release, the task switches to auditory attention, and the clock hand no longer moves but jumps to random positions without coherent motion.

### Auditory Task, Stimuli and Apparatus

When the task switches to auditory selective attention, subjects must react by pressing the button when they detect a brief silence gap that interrupts the continuous DPOAE-eliciting tones. They must focus on the auditory domain while ignoring random jumps in the position of the clock hand. Gaps occur between 1,500-2,500 ms posterior to the task switch, and 1,000 ms are given to react upon detection. Task switches to the initial passive visual period if response window ends (omitted trial) or immediately after correct detections (button press between 100-1,000 ms from gap onset), which turns the random pattern of the clock hand again into a clockwise, coherent rotation at 1 cycle per second.

Acoustic stimuli generation and recording of DPOAE were performed by RZ6 multiprocessor at a sampling rate of 48 kHz. Gaps in sound were digitally generated with squared cosine rise/fall ramps of 5 ms to avoid click-type acoustical artifacts on the onset and offset of the gap. Etymotic Research® equipment (ER10-C) specifically designed for human recording of DPOAE was used to deliver sound and record otoacoustic emissions, via three physical channels (two output phones for each primary tone, and one microphone) gently sealed within the ear canal with foam earplugs.

### Training Blocks

After we confirmed that recordings were robust (low impedance of electrophysiological signals and clear DPOAE respect to surrounding noise), we explained to subjects the instructions for the experiment, and let them practice with short blocks (11 trials) where the visual target (instantaneous clock hand position at the moment of cue offset) remained visible (as a thin line) until the response was set. We also verified that subjects heard the gaps in sound stimuli during the alternated auditory tasks, and if not, or if the visual task still was not understood, another training block was presented, with longer (easier) silence gaps if necessary (in the range 2-4 ms).

### Analysis of DPOAE Channel

For analysis of the DPOAE channel, we took a novel approach: the DPOAE amplitude was extracted from the raw microphone signal and transformed into a ‘virtual’ channel that was added to the set of electrophysiological EEG/EOG, all down-sampled to a sampling rate of 256 Hz. This allowed us to study amplitude oscillations of the cochlear amplifier in the same band of frequencies (1-35 Hz) that cognitive tasks are commonly studied in EEG research. At the technical level, this method had the purpose of taking advantage of EEG analysis techniques provided in the free software ELAN [36], for time and frequency domain measures. Two methods for taking the DPOAE amplitude were implemented by custom code in Igor 6 (Wavemetrics®). A Fourier-based method divided the signal into 16.7 ms (1/60 Hz) running windows and applied fast Fourier transform (FFT) to each. The actual length of windows was adjusted for each case, in order to be a multiple of the period of the DPOAE frequency 2f1-f2 selected for each subject. The other, a Hilbert-based method, implied first band-passing the signal through a filter with strong attenuation (>110 dB) at the frequency of the f1 primary tone, whose amplitude was typically 60-70 dB greater than the DPOAE amplitude. Having the band passed signal and its Hilbert transformation, the envelope, representing the amplitude of the DPOAE band, was calculated. The two methods yielded similar results, validating our method of continuous DPOAE amplitude extraction.

### Pre-processing of the main experiment

EEG and DPOAE artifacts were rejected by visual inspection of single trials. Rejection according to DPOAE amplitude was done by visual inspection of the complete time series from each recording block. Cursors were positioned graphically to delimit the range of valid amplitude values (see Supplementary Fig. 2 for example of DPOAE artifact removal). The unrejected DPOAE trials were then inspected one by one in the ELAN software with a selected group of channels: vertical EOG, horizontal EOG, Fourier-based DPOAE, Hilbert-based DPOAE, and then EEG channels Fz, F3, Fc1, Cz, Fc2, F4, P3, O1, Pz, O2, P4). Trials with any kind of artifact were rejected by this procedure. The remaining EEG and DPOAE trials passed to analysis with ELAN tools in the time domain and frequency domain.

### Data Analyses

Time and frequency averages across trials and all other measures were locked to either the onset of the visual cue that started the period of focused visual attention, or the attentional switch to the auditory task triggered by the behavioral response of the previous visual task (Fig. 1). For analyses purpose we used time windows of ± 1,500 ms aligned to the onset of visual or auditory attention periods.

Analysis in the time domain consisted of averaging every channel with trials locked to the onset of the visual or auditory task. In the frequency domain, we analyzed frequencies between 1 and 35 Hz, in steps of 1 Hz, with Morlet wavelets having m ratio equal to 7. First, we calculated the average of spectral z-scores. For each subject and channel, the spectrum of the single trials was obtained with the wavelet method, and frequency specific z-scores were obtained based on each trial baseline (−1,500 to 0 ms). In other words, for each trial and frequency value, the mean and standard deviation of the baseline period was calculated, and the whole spectrogram represented in z-score. Finally, these spectrograms were averaged for each subject, and then across subjects. Then, we measured the inter-trial-phase locking values, which measures the consistency of phase alignment across trials. Only the phase of each frequency component was considered and not its amplitude. Values are bounded between 0 and 1, from null to complete phase synchrony across trials.

To study the time relationship between the emergence of cortical and cochlear oscillations revealed by the average of spectral z-scores, we measured for each subject the time points where 50% of the maximum oscillatory power was achieved in the band between 1 and 7 Hz, for Fz, Cz, O2 and the DPOAE-amplitude channels. These time values are box-plotted in Fig 4c,d, and the statistical significance between channels evaluated with Mann-Whitney (using p<0.05 as significant).

## Acknowledgements

We thank Jean-Pierre Aguera for his kind support regarding the use of ELAN, an open software for EEG analysis and Conrado Bosman for his critical review of a previous version of this manuscript. Funded by Proyecto Fondecyt 1161155 to P.H.D. and Fundacion Guillermo Puelma.

## Author Contributions

**Conceptualization:** Paul H. Delano, Constantino Dragicevic, Luis Robles.

**Data curation:** Constantino Dragicevic, Bruno Marcenaro, Marcela Navarrete.

**Formal analysis:** Constantino Dragicevic.

**Funding acquisition:** Paul H. Delano.

**Investigation:** Constantino Dragicevic, Bruno Marcenaro, Marcela Navarrete.

**Methodology:** Constantino Dragicevic, Bruno Marcenaro, Marcela Navarrete.

**Project administration:** Paul H. Delano.

**Software:** Constantino Dragicevic, Bruno Marcenaro.

**Supervision:** Paul H. Delano, Luis Robles.

**Visualization:** Constantino Dragicevic, Bruno Marcenaro, Marcela Navarrete, Paul H. Delano, Luis Robles.

**Writing – original draft:** Constantino Dragicevic.

**Writing – review and editing:** Constantino Dragicevic, Bruno Marcenaro, Marcela Navarrete, Paul H. Delano, Luis Robles.

## Figure captions

**Supplementary Figure 1. DPOAE band spectra of individual subjects.** FFT of the band-pass filtered microphone signal of each subject prior to Hilbert transform. Each subplot corresponds to one block of one subject. Filters were centered at DPOAE frequency (2 f1 – f2), and had a flat frequency response with no attenuation in a ±50 Hz vicinity. Amplitude is shown in attenuation dB relative to the DPOAE peak amplitude.

**Supplementary Figure 2. DPOAE-based artifact rejection.** Representative example of the time course of DPOAE amplitude in one block of a subject, illustrating the first rejection stage based solely on DPOAEs. Horizontal dashed lines were manually positioned to define rejection limits (shown by “A” and “B”). Only trials unrejected by this process were further scrutinized in the following EEG artifact rejection stage.

